# Modelling within-host macrophage dynamics in influenza virus infection

**DOI:** 10.1101/2020.05.07.083360

**Authors:** Ke Li, James M. McCaw, Pengxing Cao

## Abstract

Human respiratory disease associated with influenza virus infection is of significant public health concern. Macrophages, as part of the front line of host innate cellular defence, have been shown to play an important role in controlling viral replication. However, fatal outcomes of infection, as evidenced in patients infected with highly pathogenic viral strains, are often associated with prompt activation and excessive accumulation of macrophages. Activated macrophages can produce a large amount of pro-inflammatory cytokines, which leads to severe symptoms and at times death. However, the mechanism for rapid activation and excessive accumulation of macrophages during infection remains unclear. It has been suggested that the phenomena may arise from complex interactions between macrophages and influenza virus. In this work, we develop a novel mathematical model to study the relationship between the level of macrophage activation and the level of viral shedding in influenza virus infection. Our model combines a dynamic model of viral infection, a dynamic model of macrophages and the essential interactions between the virus and macrophages. Our model predicts that the level of macrophage activation can be negatively correlated with the level of viral shedding when viral infectivity is sufficiently high. We further identify that temporary depletion of resting macrophages in response to viral infection is a major driver in our model for the negative relationship between macrophage activation and viral shedding, providing new insight into the mechanisms that regulate macrophage activation. Our model serves as a framework to study the complex dynamics of virus-macrophage interactions and provides a mechanistic explanation for existing experimental observations, contributing to an enhanced understanding of the role of macrophages in influenza viral infection.

## 1. Introduction

Influenza is a contagious respiratory disease caused by influenza viruses. Infection with influenza A virus (IAV) in particular remains as a major public health concern, resulting in heavy morbidity worldwide every year [1]. Epithelial cells, which line the upper respiratory tract (URT) of the host, are the primary target cells for influenza virus infection [2, 3], and virus-induced cell damage is often thought to be the main cause for clinical symptoms and a determinant of virulence [4, 5, 6, 7]. During an infection, host immunity plays an important role for viral resolution and host recovery. The innate (or nonspecific) immune system is the first and primary defence mechanism that is triggered upon detection of an IAV infection. Macrophages, as part of the innate immune cellular response, are activated at the early stages of infection [8, 9, 10]. They perform two important antiviral functions. One is the uptake of viruses mediated by the interaction of pattern-recognition receptors (PRRs), such as the Toll-like receptors (TLRs), in macrophages with pathogen-associated molecular patterns (PAMPs) on the virus, and the phagocytosis of apoptotic virus-infected cells [11, 12, 13, 14]. The other is the secretion of cytokines and chemokines, such as tumor necrosis factor-alpha (TNF-*α*), interleukins-6 (IL-6) and interferons (IFNs), by which macrophages can modulate inflammatory responses and help trigger an adaptive immune response, attracting effector cells to the site of infection. [15, 16, 17].

Macrophages are highly heterogeneous in the host and can alter their phenotypes and functions rapidly in response to local stimuli [18, 19, 20]. In response to a viral infection, resting macrophages are activated and give rise to two major types of macrophages, denoted *M*_1_ and *M*_2_ in terms of functionality [21]. *M*_1_ macrophages have a stronger capability to engulf free virions, present antigens to other immune cells and produce pro-inflammatory cytokines which contribute to both host inflammatory responses and pathogen clearance [18]. In contrast, *M*_2_ macrophages primarily secrete anti-inflammatory cytokines to mitigate inflammation and maintain host homeostasis [22]. While viral infection-induced cell death and tissue damage are thought to be the primary contributors to host morbidity, further evidence has shown that over-expression of pro-inflammatory cytokines and chemokines mediated by activated macrophages may also be a cause for lung pathology [23, 24, 25, 26, 24, 27]. Unregulated pulmonary infiltration of macrophages is often the hallmark of severe influenza virus infection [28, 29, 30], as reviewed in [31]. For instance, mice infected with highly pathogenic (HP) influenza virus experienced rapid infiltration of macrophages and excessive accumulation of macrophages during infection, showing fatal infection results with a high level of viral shedding. These outcomes were not observed in mice infected with low virulent strains [32]. The observations suggest a positive correlation between the level of macrophage activation and the level of viral shedding. However, since the interactions between different types of macrophages and between macrophages and influenza virus involve both positive and negative feedback mechanisms (see review [33]), it is not clear how this relationship can arise from a dynamical system of virus-macrophage interactions and under what condition(s) such a relationship may no longer be valid. In this paper, we study these interactions using a mathematical model.

Mathematical models have been used to explore macrophage dynamics in different pathological environments, e.g., in bacterial infection [34, 35] (particularly in tuberculosis infection (TB) [36]) and for tumors [37]. Some models in the literature have been used to specifically investigate the interactions between *M*_1_ and *M*_2_ macrophages [38, 39]. Models have also been used to study within-host influenza virus dynamics, many of which have been designed to explore how the viral load kinetics is modulated by different immunological factors (see reviews [40, 41, 42]). However, in the literature, there are no influenza virus infection models which explicitly include both *M*_1_ and *M*_2_ macrophage dynamics as part of the innate immune responses.

In this paper, we develop a novel mathematical model, which combines a dynamic model of influenza virus infection, a dynamic model of macrophages and the essential interactions between virus and macrophages. We use the model to explore the dynamics of macrophages in response to influenza viral infection and investigate how the level of macrophage activation is influenced by the viral infectivity (which is a critical parameter determining the level of viral shedding). Our aim is to explore in detail possible explanation(s) for the mechanism determining the aforementioned relationship between viral shedding and macrophage activation. Finally we discuss our findings and the biological implications of our model results.

## 2. Methods

We first introduce a dynamic model of macrophages, which incorporates three distinct populations and the conversion processes between them, in the absence of viral infection. We then incorporate the macrophage model into an influenza viral infection model which captures the minimal essential processes to describe IAV kinetics, including viral multiplication via the infection of epithelial cells and viral resolution by antibodies.

### 2.1. A dynamical model of macrophages

The model of macrophage dynamics contains three macrophage populations: *M*_1_ (so called “classically activated” macrophages), *M*_2_ (so called “alternatively activated” macrophages) [21] and resting macrophages *M* which have low efficiency to present antigen and produce cytokines. Resting macrophages can convert into either *M*_1_ or *M*_2_ macrophages in response to different stimuli [21, 36]. A diagram representing the model is shown in Fig. 1.

**Figure 1:**
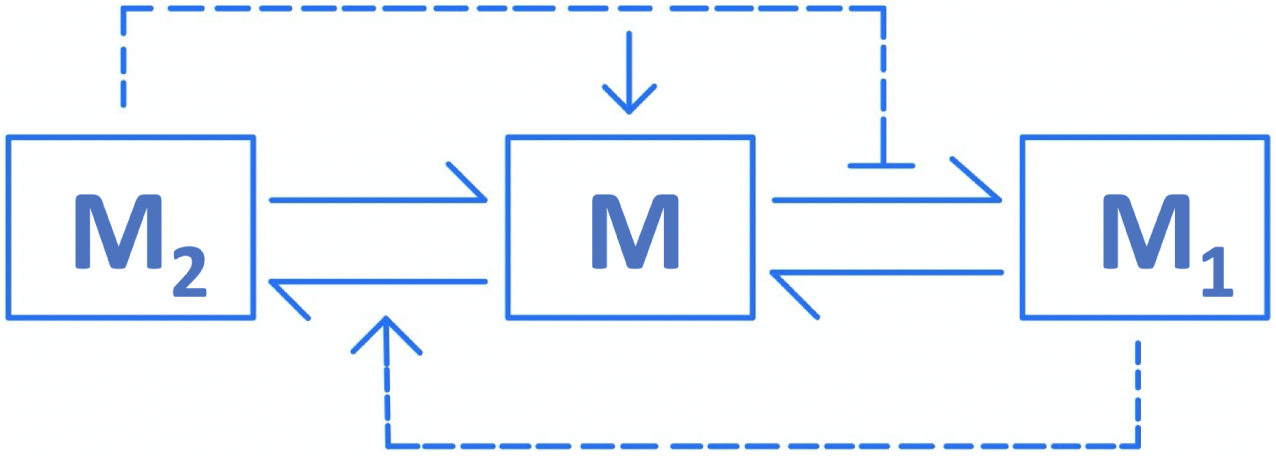
Macrophage dynamics in the absence of viral infection. The dashed arrow line indicates that the presence of *M*_1_ promotes the conversion process of *M* to *M*_2_. The dashed-bar line denotes suppression of the process of *M* to *M*_1_ due to *M*_2_. The solid full-arrow line denotes recruitment of *M*, and solid half-arrow lines denote the conversion processes among macrophages.

In the absence of infection, *M* macrophages can be converted into *M*_1_ in response to apoptotic cells as well as tissue debris, and *M*_1_ macrophages stimulate the secretion of pro-inflammatory cytokines, e.g. TNF-*α* and IFN- *γ* [43], which subsequently reinforce the activation process in an autocrine or paracrine manner [44]. *M* macrophages can also be converted into *M*_2_ macrophages stimulating the secretion of anti-inflammatory cytokines, e.g. interleukin-10 (IL-10), to mitigate the host inflammatory response and maintain homeostasis [19]. The conversion processes are modelled by a set of ordinary differential equations (ODEs):

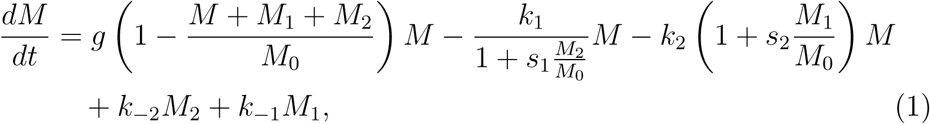

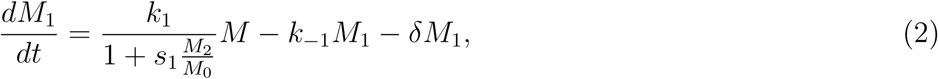

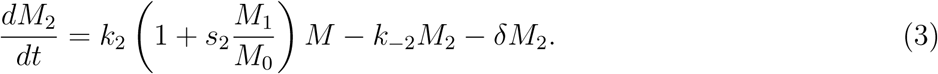

Eq. 1 describes the rate of change of resting macrophages *M*. It is governed by five processes (corresponding to the five terms on the righthand side of Eq. 1). *M* are produced at a rate *g*(1 − (*M* + *M*_1_ + *M*_2_)*/M*_0_)*M* mimicking a logistic growth model. To phenomenologically capture the established regulatory effects of *M*_1_ and *M*_2_ [23] as reviewed in [21], the rate of conversion from *M* to *M*_1_ is modelled by a decreasing function of *M*_2_ with a maximum of *k*_1_ (see the second term on the righthand side of Eq. 1) and the rate of conversion from *M* to *M*_2_ is modelled by an increasing function of *M*_1_ with a minimum of *k*_2_ (see the third term on the righthand side of Eq. 1). The parameters *s*_1_ and *s*_2_ modulate the dependence of the conversion rates on the number of activated macrophages. *M*_1_ and *M*_2_ macrophages return to the resting state *M* at rate *k*_−1_ and *k*_−2_ when stimuli are diminished, respectively [45].

Eq. 2 and Eq. 3 model the dynamics of *M*_1_ and *M*_2_, respectively. In addition to the terms describing the conversion between *M* and *M*_1_ (i.e. the first and second terms on the righthand side of Eq. 2) and between *M* and *M*_2_ (i.e. the first and second terms on the righthand side of Eq. 3), macrophages *M*_1_ and *M*_2_ decay naturally at rate *δ*.

### 2.2. A model coupling macrophage dynamics and viral infection dynamics

Upon detection of virus, resting macrophages *M* are promptly activated and converted into *M*_1_ via Toll-like receptor (TLR)-dependent signalling pathways, and strong inflammatory responses are initiated [14]. A wide range of inflammatory cytokines and chemokines, such as tumor necrosis factor-alpha (TNF-*α*), interleukins-6 (IL-6) and interferons (IFNs), are secreted by proinflammatory macrophages *M*_1_ and lead to the recruitment of more immune cells to the site of infection [46]. Activated macrophages have been shown to have a stronger capacity for phagocytosis of apoptotic cells and antigen presentation compared to those in an inactivated state [47].

Macrophages are activated in response to viral infection and are crucial in providing negative feedback to viral reproduction. For example, activated *M*_1_ macrophages uptake free virions and prime various adaptive immune responses [16, 21]. Here we propose a model to capture the essential regulatory processes between macrophages and influenza virus. A diagram representing the model is shown in Fig. 2, and the processes are modelled by a system of ODEs:

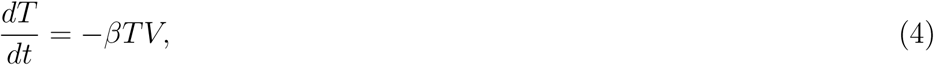

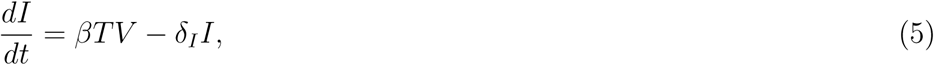

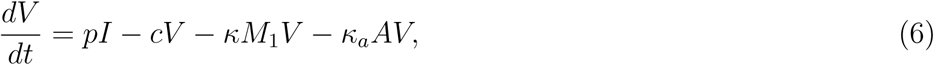

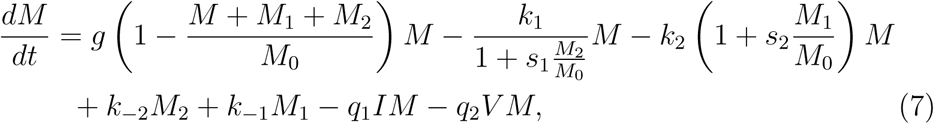

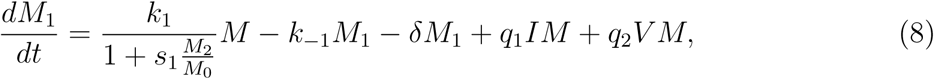

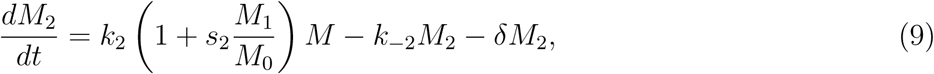

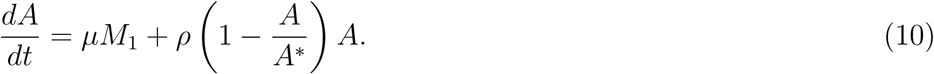

**Figure 2:**
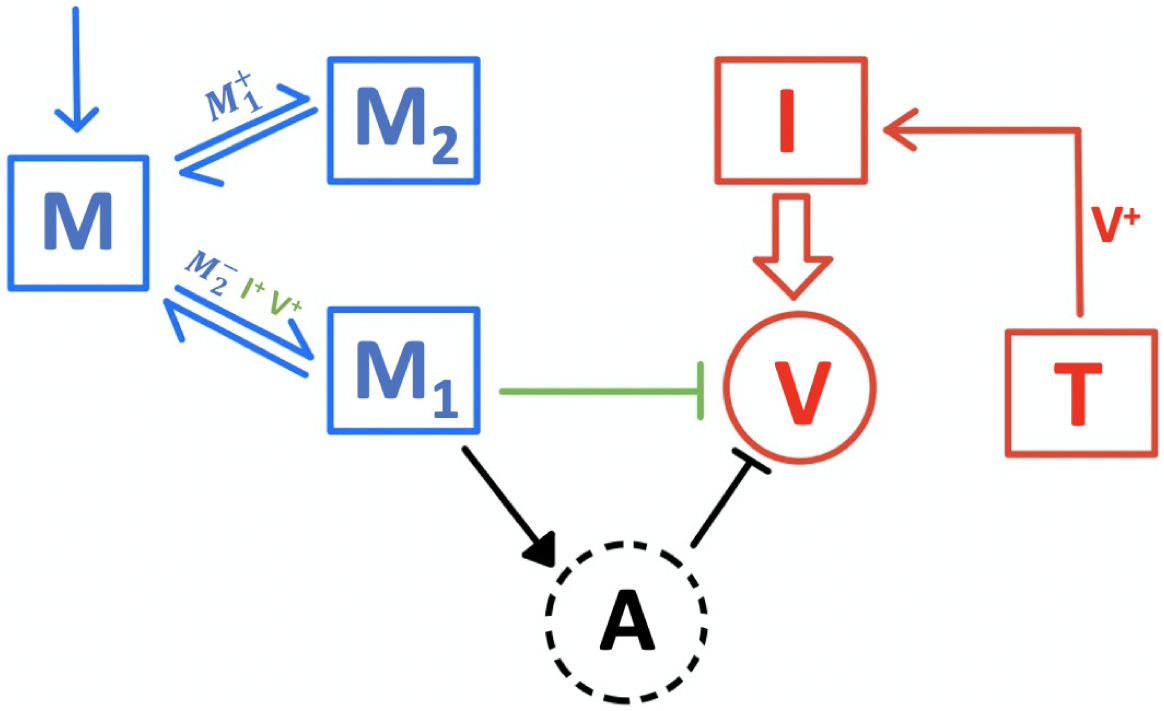
Macrophage dynamics in the presence of viral infection. Resting macrophages (*M*) replenish their population in the system (blue arrow towards *M*) and are activated into either pro-inflammatory macrophages *M*_1_, or anti-inflammatory macrophages *M*_2_ (blue arrows from *M* to either *M*_1_ or *M*_2_). The activated macrophages return back to *M* (as indicated by the blue arrows from *M*_1_/*M*_2_ to *M*). 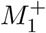 indicates that *M*_1_ increases the rate of conversion from *M* to *M*_1_. In a similar way, 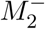, *V* ^+^ and *I*^+^ indicate that the rate of conversion from *M* to *M*_1_ is suppressed by *M*_2_ but enhanced by virus *V* and infected cells *I*. Virus (*V*) infects epithelial cells (*T*), which become infected cells (*I*) (indicated by the red solid arrow with *V* ^+^), and infected cells (*I*) produce virus (large arrow). *M*_1_ internalise free virions (green solid bar line), and stimulate adaptive immunity (black solid full-triangle line) in which antibodies (*A*) are produced, by which virions are neutralised (black solid bar line).

Eqs. 4–6, proposed based on the classic target cell-infected cell-virus (TIV) model, describe the essential dynamics of virus turnover through the infection of target cells and the resolution of infection by immune responses. In detail, target cells (*T* ; i.e. epithelial cells in influenza infection) are infected with virus (*V*) and become infected cells (*I*) at a rate *βV*. Infected cells produce and release viral progenies (at a rate *p*) which invade target cells leading to further infection. Free virus (*V*) decays due to three processes: natural decay at a rate *c*, internalisation by activated macrophages *M*_1_ at a rate *κM*_1_ and neutralisation by antibodies at a rate *κ*_*a*_*A*. Infected cells (*I*) die naturally at a rate *δ*_*I*_.

Eqs. 7–9 are adapted from the macrophage model (Eqs. 1–3) with some additional terms capturing the effect of infected cells and virus on the conversions from *M* to *M*_1_ and *M*_2_. For example, the term *q*_1_*IM* models the conversion of resting macrophages *M* to *M*_1_ due to the presence of various infected cell-producing cytokines [48, 49]. The term *q*_2_*V M* models virus-induced macrophage activation via TLR-dependent pathways [16].

Eq. 10 models the activation and expansion of adaptive immune responses, in particular the production of antibodies, which are responsible for clearing virus at the late stages of infection, providing long-term protection. In detail, the production of antibodies (*A*) is phenomenologically modelled by a logistic growth model (i.e. the second term on the righthand side of Eq. 10) with a growth rate *ρ* and a carrying capacity *A*^*\**^, coupled with a “trigger” term due to antigen presentation, *µM*, which assumes that the strength of triggering the adaptive immune response is proportional to the level of activated macrophage *M*_1_.

### 2.3. Model parameters

The values of model parameters are given in Table 1. The parameter values and initial conditions for influenza viral dynamics (such as *p, δ*_*I*_, *T* (0), *I*(0) and *V* (0)) are chosen from the study in [50], in which the authors fitted the TIV model to a set of data from humans infected with A/H1N1 virus. The parameter *β* is estimated and chosen from the literature such that the viral load shows at least a three-fold increase in infection [51]. To the best of our knowledge, the values for the model parameters which govern either the macrophage dynamics (such as *M*_0_, *k*_1_, *k*_−1_, *k*_2_ and *k*_−2_) or the influenza virus-macrophage interactions (such as *q*_1_ and *q*_2_), are not available. Therefore, we choose those parameter values from [52] in which macrophage dynamics are investigated in a tumour environment. We assume macrophages (*M*_1_ and *M*_2_) have comparable impact on each other and set *s*_1_ = *s*_2_ = 1. Also, the assumed carrying capacity for antibodies (*A*^*\**^) during infection is chosen such that virus can be efficiently cleared.

**Table 1:** Parameter values used for numerical simulation. [·] denotes the unit for each variable, e.g., the unit of *T* is denoted as [*u*_*T*_]. *d*^−1^ denotes per day.

| Par. | Description | Value | Unit | Reference |
| --- | --- | --- | --- | --- |
| $p$ | Viral production rate | $7.1 \times 10^{-2}$ | $[u_V][u_T]^{-1}d^{-1}$ | [50] |
| $c$ | Viral natural death rate | 20 | $d^{-1}$ | [51] |
| $s_1$ | Effectiveness of $M_2$ attenuates $M \rightarrow M_1$ | 1 | - | - |
| $s_2$ | Effectiveness of $M_1$ promotes $M \rightarrow M_2$ | 1 | - | - |
| $\delta_I$ | Natural death rate of infected cells | 3.6 | $d^{-1}$ | [50] |
| $\delta$ | Decay rate of $M_1$ and $M_2$ | 0.02 | $d^{-1}$ | [52] |
| $\kappa$ | Rate of virus internalisation by $M_1$ | $7.7 \times 10^{-4}$ | $[u_{M_1}]^{-1}d^{-1}$ | [34] |
| $\kappa_a$ | Neutralisation rate of virus by antibody | 0.2 | $[u_A]^{-1}d^{-1}$ | [51] |
| $\mu$ | Rate of macrophage-induced activation of adaptive immunity | $10^{-6}$ | $d^{-1}$ | - |
| $\rho$ | logistic growth rate of antibody response | 1 | $d^{-1}$ | [53] |
| $\beta$ | Viral infectivity | $3.8 \times 10^{-5}$ | $[u_V]^{-1}d^{-1}$ | - |
| $g$ | Regrowth rate of $M$ | 0.2 | $d^{-1}$ | - |
| $M_0$ | Carrying capacity of macrophage regrowth | $10^6$ | $[u_M]$ | [34, 37] |
| $k_1$ | Conversion rate of $M \rightarrow M_1$ | 0.5 | $d^{-1}$ | [52] |
| $k_{-1}$ | Conversion rate of $M_1 \rightarrow M$ | 0.33 | $d^{-1}$ | [52] |
| $k_2$ | Conversion rate of $M \rightarrow M_2$ | 0.5 | $d^{-1}$ | [52] |
| $k_{-2}$ | Conversion rate of $M_2 \rightarrow M$ | 0.33 | $d^{-1}$ | [52] |
| $q_1$ | Rate of infected cell-induced conversion from $M$ to $M_1$ | $1 \times 10^{-6}$ | $[u_I]^{-1}d^{-1}$ | [34] |
| $q_2$ | Rate of virus-induced conversion from $M$ to $M_1$ | $1 \times 10^{-6}$ | $[u_V]^{-1}d^{-1}$ | [34] |
| $A^*$ | Assumed carrying capacity of antibody upon an infection | $10^5$ | $[u_A]$ | - |
| $T(0)$ | Initial value of uninfected epithelial cells in the upper respiratory tract | $4 \times 10^8$ | $[u_T]$ | [50] |
| $I(0)$ | Initial value of infected cells | 0 | $[u_I]$ | [50] |
| $V(0)$ | Initial value of viral load | $3.3 \times 10^{-1}$ | $[u_V]$ | [50] |

### 2.4. Numerical simulation methods

The ordinary differential equations are solved using the ode solver *ode15s* in MATLAB R2019b with a relative tolerance of 1 *×* 10^−5^ and an absolute tolerance of 1 *×* 10^−10^. The initial values for the target cell *T*, the infected cell *I* and the virus *V* are given in Table 1. The initial values for the macrophage populations *M, M*_1_ and *M*_2_ are given by the virus-free steady state and are obtained by numerically integrating Eqs. 1–3 (using *ode15s* in MATLAB) for a sufficiently long time interval (see Fig. S1), and choosing the values of *M, M*_1_ and *M*_2_ at the final time point. MATLAB code to produce all the figures in this study can be found at https://github.com/keli5734/Matlab-Code.

## 3. Results

### 3.1. Dynamics of macrophages and viral shedding

The simulated time series of viral load (*V*) and two populations of activated macrophages (*M*_1_ and *M*_2_) are shown in Fig. 3A (model solutions for other variables are given in Fig. S2 in the *Supplementary Material 1*). The viral load curve shows a three-phase shape—an exponential growth followed by a slow decay (or a plateau) and finally a rapid decay to viral resolution— typical of observed infection data and simulations of *in vivo* influenza infection [50, 51, 53, 54, 55]. In response to viral infection, activated macrophages *M*_1_ undergo a rapid increase followed by a decrease (Fig. 3A red solid line), while *M*_2_ macrophages experience a decline followed by a replenishment (Fig. 3A red dashed line). The different behaviours of *M*_1_ and *M*_2_ are due to competition for the limited resource (i.e. resting macrophage *M*). There is a dramatic increase in the conversion from *M* to *M*_1_ induced by the initial exponential viral growth that rapidly consumes *M* (see Fig. S2) which in turn reduces the conversion from *M* to *M*_2_. *M*_1_ and *M*_2_ gradually return to their homeostatic state upon the resolution of infection (after approximately day 12 in Fig. 3A).

**Figure 3:**
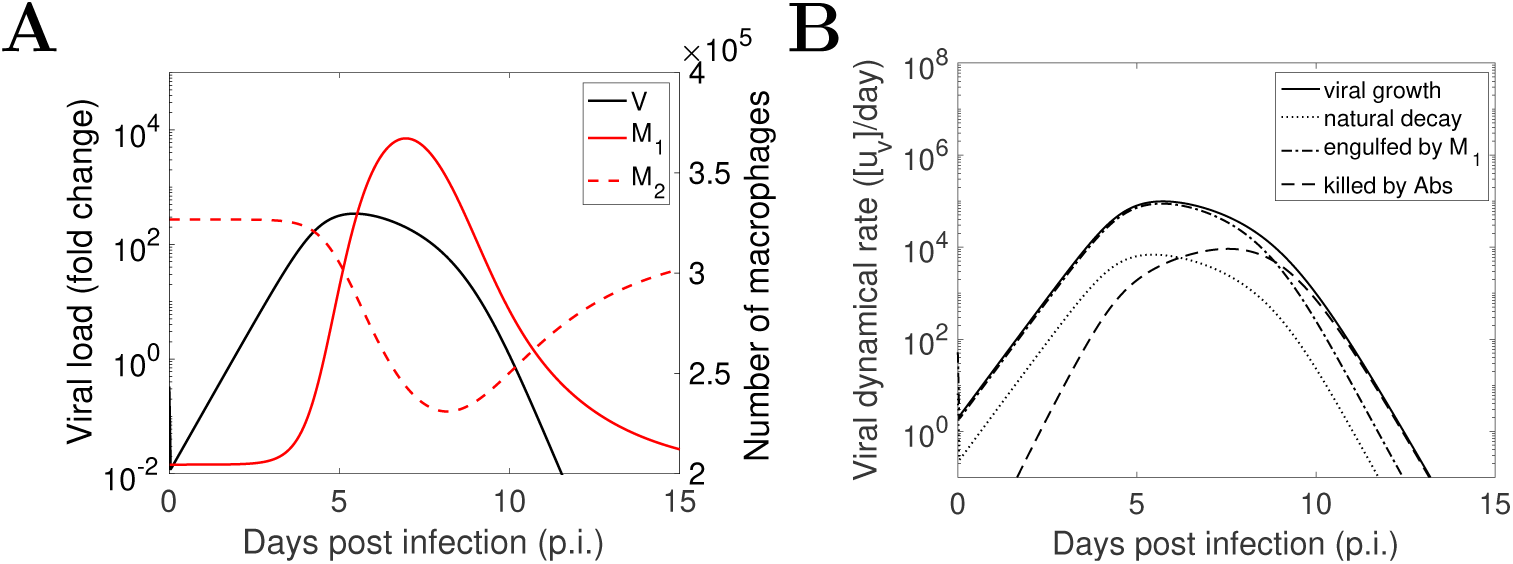
Model simulation results of viral shedding kinetics and macrophage dynamics in infection using the parameters values in Table 1. (A) Viral shedding dynamics (Eq. 6; black solid line), pro-inflammatory macrophages *M*_1_ (Eq. 8; red solid line) and anti-inflammatory macrophages *M*_2_ (Eq. 9; red dashed line). (B) The rate of change of the components on the right-hand side of *dV/dt* (Eq. 6), which are the rates of viral growth *pI* (solid line), viral natural decay *cV* (dot line), virus engulfed by *M*_1_ macrophages *κM*_1_*V* (dash-dotted line), and virus neutralized by antibodies *κ*_*a*_*AV* (dashed line).

To better understand how the viral load is influenced by macrophages, we present the time series of the four terms on the righthand side of Eq. 6 (Fig. 3B). These four time-series represent the four major processes determining the rate change of viral load (*dV/dt*) in the model and include viral production (*pI*), natural death (*cV*), internalisation by *M*_1_ macrophage (*κM*_1_*V*) and neutralisation by antibodies (*κ*_*a*_*AV*). The macrophage-mediated innate immune response (dash-dotted line) plays a dominant role in controlling viral replication before antibody takes over on approximately day 9 in the model. A qualitatively similar model behaviour was observed in [51] where the innate immune response was assumed to be mediated by interferon.

### 3.2. Relationship between the level of macrophage activation and the level of viral shedding

Having examined the time-series behaviour of viral shedding and *M*_1_ macrophages, we now examine the relationship between the level of macrophage activation and the level of viral shedding. The level of macrophage activation is assumed to be the cumulative number of *M*_1_ macrophages because of their key role in producing massive pro-inflammatory cytokines in influenza virus infection [56], and we quantify the cumulative number by the area under the *M*_1_ time-series curve (AUC_*M*1_)

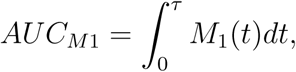

where *τ* is a cut-off day for computation. The level of viral shedding is assumed in the model to be the cumulative viral load, which has been considered as a surrogate for viral infectiousness of influenza infection [57] and an important marker for viral pathogenicity [58, 59]. It is quantified by the area under the viral load time-series curve (AUC_*V*_)

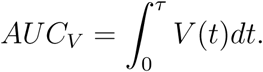

In this study, we set *τ* = 15 which is an appropriate value to cover both the duration of viral infection and the duration of macrophage activation as shown in [32].

Fig. 4 shows the relationship between AUC_*M*1_ and the AUC_*V*_ as viral infectivity (model parameter *β*) varies. We chose to vary the viral infectivity because it is a key parameter determining the ability of virus to cause infection. We see that for intermediate values of *β*, the AUC_*M*1_ and the AUC_*V*_ are positively correlated (e.g. in Region II; the definition of the regions are provided in the caption of Fig. 4), consistent with experimental observation [32]. However, we also identify in the model a region where the two quantities are negatively correlated for relatively high *β* (i.e. Region III), which suggests that a highly pathogenic virus strain may cause a compromised activation of *M*_1_ macrophages while maintaining a high level of viral shedding. In addition, for very small *β* (i.e. in Region I), viral infection cannot be established because the basic viral reproduction number (provided in the caption of Fig. 4) is less than the infection threshold 1.

**Figure 4:**
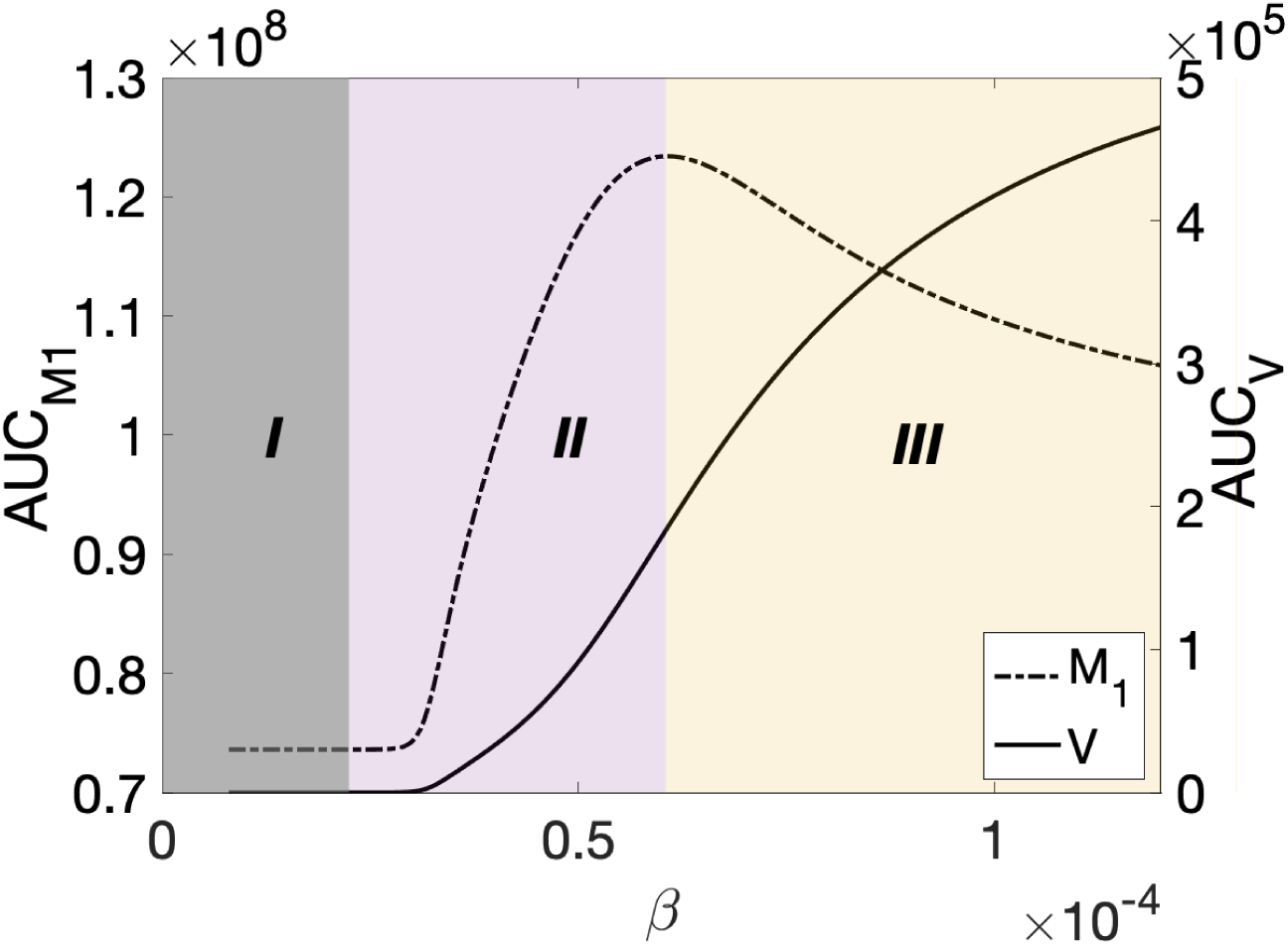
Simulation results of the change of AUC_*M*1_ (dash-dotted line) and AUC_*V*_ (solid line) to the modulation of viral infectivity. *β ∈* [8 *×* 10^−6^, 1.2 *×* 10^−4^]. We classify the results into three regions. Regions I and II are separated by the viral basic reproduction number which is given by *R*_*V*,0_ = *pβT* (0)*/*(*δ*_*I*_ (*c* + *κM*_1_(0) + *κ*_*a*_*A*(0))). In region I, *R*_*V*,0_ *<* 1. At the boundary between regions I and II, *R*_*V*,0_ = 1. In regions II and III, *R*_*V*,0_ *>* 1. The boundary between regions II and III is determined by a change in the correlation between AUC_*M*1_ and AUC_*V*_. In region II these two areas are positively correlated whereas in region III they are negatively correlated.

It is unclear from Fig. 4 why the negative relationship between the AUC_*M*1_ and the AUC_*V*_ in region III arises, so we examine the time series for the viral load and *M*_1_ macrophages. Fig. 5A and 5B show the time series of viral load and different types macrophages for *β* = 6.08 *×* 10^−5^ which is the critical value separating regions II and III. Fig. 5C and 5D show similar time series for a *β* value inside the region III (i.e. *β* = 10.08 *×* 10^−5^). Although the viral load curve for the larger *β* exhibits a shorter duration of infection compared to that for the smaller *β* (Fig. 5C v.s. Fig. 5A), it also exhibits a higher peak value such that a higher AUC_*V*_ is possible. In contrast, the *M*_1_ macrophage curve for the larger *β* exhibits both a lower peak value and a shorter duration of activation—quickly reaching a peak and declining—compared to that for the smaller *β* (Fig. 5D v.s. Fig. 5B), which explains the decrease in AUC_*M*1_. In the next section, we will explore the mechanism(s) leading to the reduction in AUC_*M*1_.

**Figure 5:**
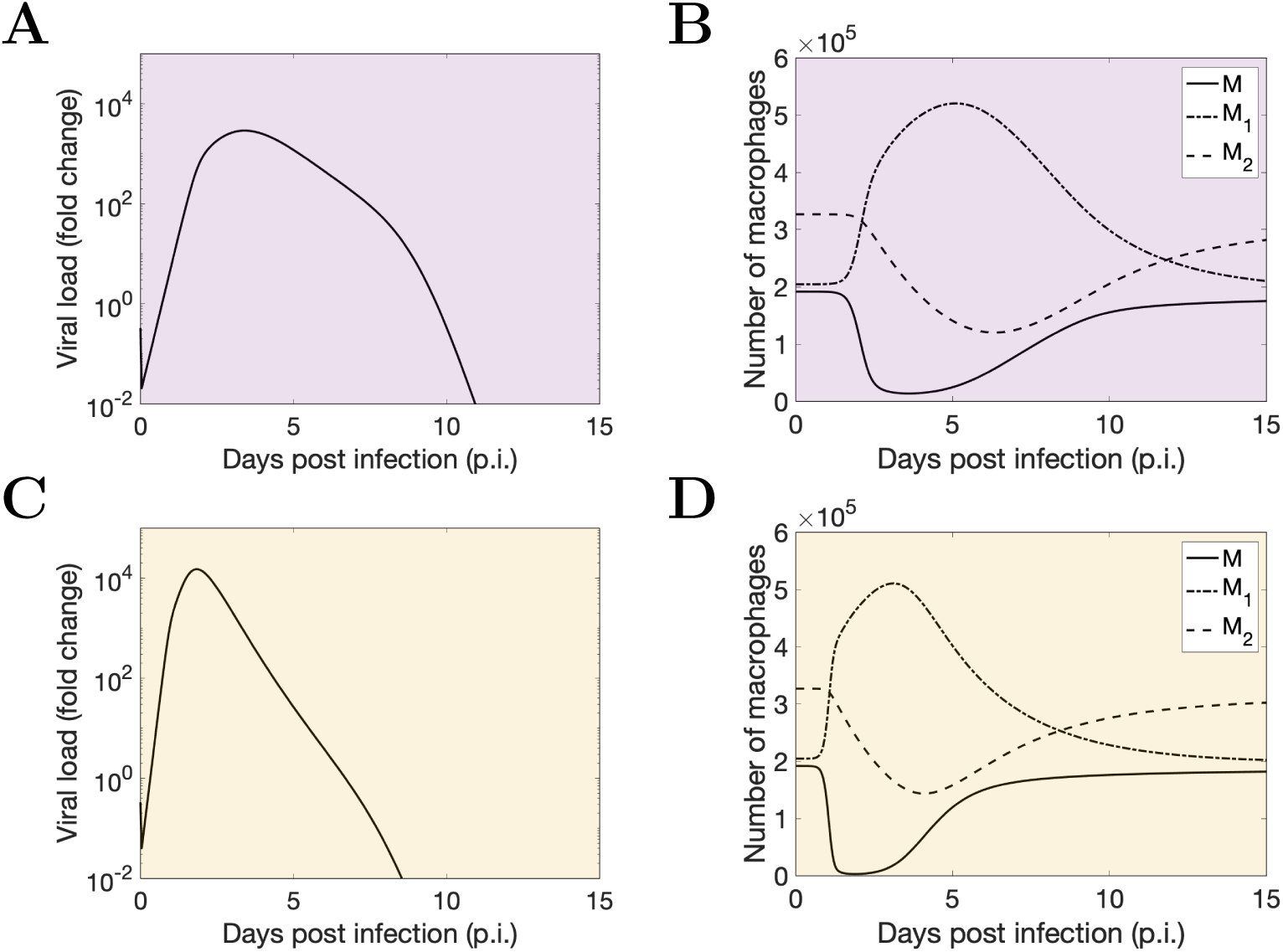
Model simulation results of macrophage dynamics and viral shedding kinetics with different viral infectivity *β* in different regions. First row (A and B): viral shedding kinetics, and the dynamics of *M* macrophages (solid line), *M*_1_ macrophages (dash-dotted line) and *M*_2_ macrophages (dashed line) in region II (*β* = 6.08 *×* 10^−5^). Second row (C and D): viral shedding kinetics and macrophage dynamics in region III (*β* = 10.08 *×* 10^−5^).

### 3.3. Temporary depletion of M is a mechanism driving the decrease of AUC_M1_ in region III

Since the production of *M*_1_ macrophages is fundamentally driven by the conversion of resting macrophages *M* in the model (shown in Fig. 1), we hypothesise that the decrease in AUC_*M*1_ in region III might be attributed to a more severe (albeit temporary) depletion of *M*, which is partly supported by Fig. 5B and 5D where the level of *M* is driven lower and earlier for a larger *β*.

To investigate our hypothesis, we increase the regrowth rate of *M* (i.e. the model parameter *g*), which should mitigate the extent of *M* depletion by increasing both the rate of *M* replenishment and the initial number of *M* macrophages (i.e. the homeostatic state, see Fig. 6A). As seen in Fig. 6B, as *g* increases, region III shrinks while both region I and region II expand. Fig. 6C provides a more detailed view of how the regions shift for two selected values of *g* (in particular the expansion of region II at the expense of a shrinking region III). These results show that mitigating *M* depletion (by increasing the regrowth rate *g* of *M* macrophages in the model) can turn a decreasing AUC_*M*1_ to increasing for a range of *β*, confirming our hypothesis that a temporary depletion of *M* is a mechanism driving the decrease of AUC_*M*1_ in region III.

**Figure 6:**
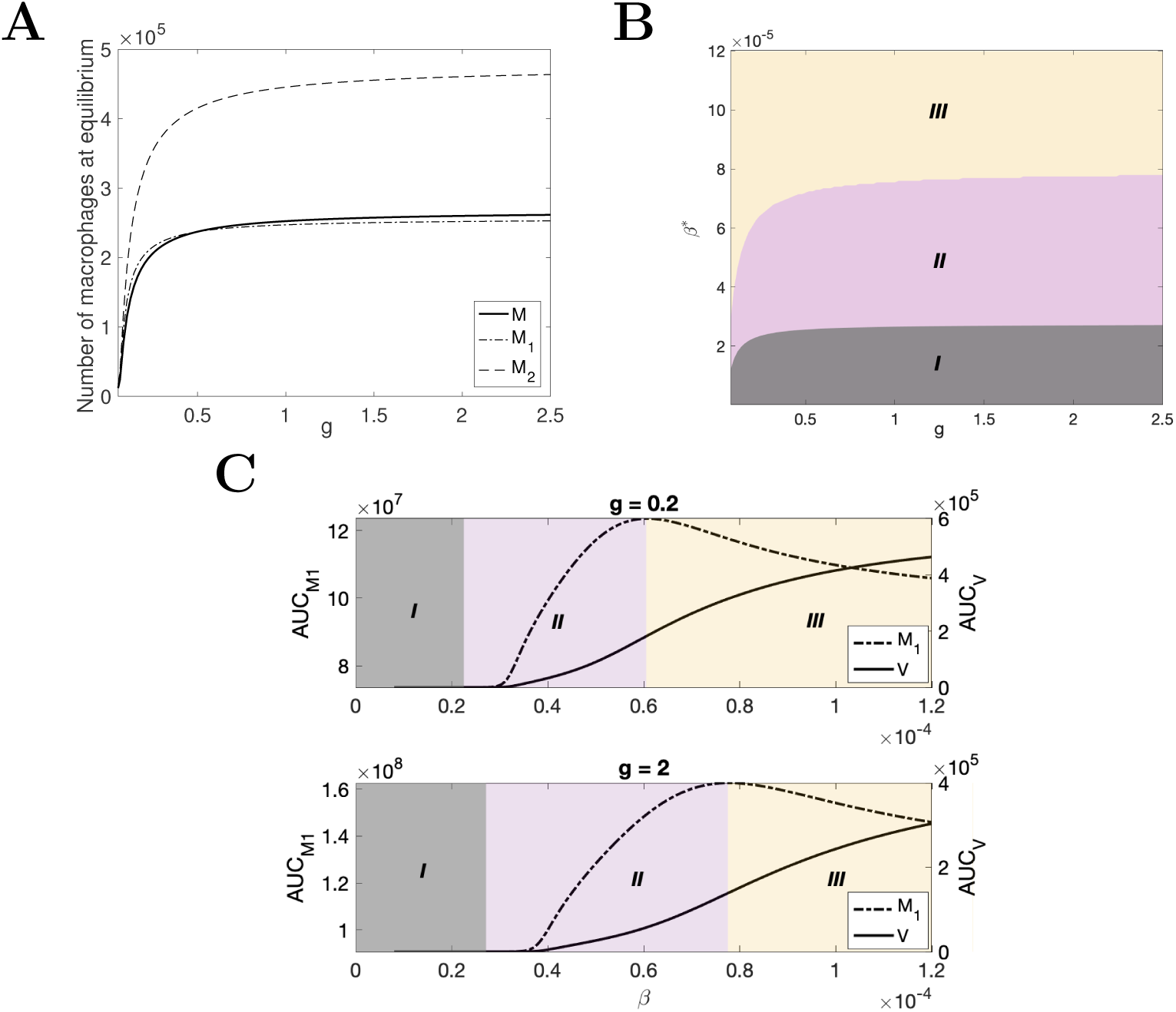
Model simulation results of macrophages, AUC_*M*1_ (dashed line) and AUC_*V*_ (solid line) as functions of *M* regrowth rate *g*. (A) The dependence of initial values of macrophage populations on the *M* regrowth rate *g. g* is varied from 0.02 to 2.5. Note that the numbers of macrophages become saturated for relatively large *g*. We show mathematically that there always exists a maximum capacity for *M* as *g* → *∞* (see *Supplementary Materials 2* for detail). (B) shows how regions I, II and III change as *g* increases. (C) The dependence of the AUC_*M*1_ and the AUC_*V*_ on *β* for two selected values of *g* (i.e. *g* = 0.2 and *g* = 2). *β* ∈ [8 *×* 10^−6^, 1.2 *×* 10^−4^].

### 3.4. Dependence of the AUC_V_ -AUC_M1_ relationship on other model parameters

So far we have varied the viral infectivity *β*, as a means to examine the relationship between the level of viral shedding and the level of *M*_1_ macrophage activation. We now examine whether our results and conclusions are robust to a change in other virus-related parameters. For example, we vary the viral production rate *p* and produce a series of figures similar to Figs. 4–6 (see Figs. S3–S5 in *Supplementary Material 1*). We find that the results are qualitatively the same as those for varying *β*, in particular the existence of region III (see Fig. S3) and the observed reduction in region III when mitigating depletion of *M* by increasing the regrowth rate *g* (see Fig. S5). Varying *κ* (the rate of viral engulfment by *M*_1_) yields similar results (see Figs. S6–S8; note that the effect of decreasing *κ* is similar to that of increasing *β* or *p* because of the antagonistic processes described by the model).

Since experimental studies [26, 27, 29, 30] have suggested highly pathogenic influenza virus strains can infect macrophages and modulate the rate of cytokine production, resulting in rapid macrophage infiltration and strong inflammatory response, we further examine the effect of the rate of *M*_1_ activation, either virus-induced (*q*_1_) or virus-independent (*k*_1_), on the AUC_*V*_ – AUC_*M*1_ relationship. Fig. 7A shows that increasing *q*_1_ leads to an increase in AUC_*M*1_ but a decrease in AUC_*V*_. A similar result is observed for increasing *k*_1_ (Fig. 7B). The negative relationship between AUC_*V*_ and AUC_*M*1_ in response to a change in the rate of *M*_1_ activation in the model may imply that a viral mutation conveying a change in the rate of *M*_1_ activation will result in either compromised viral shedding or compromised macrophage response and therefore cannot solely explain the aforementioned observation that highly pathogenic influenza virus strains can induce both a higher cell infiltration and a higher level of viral shedding than less pathogenic strains.

**Figure 7:**
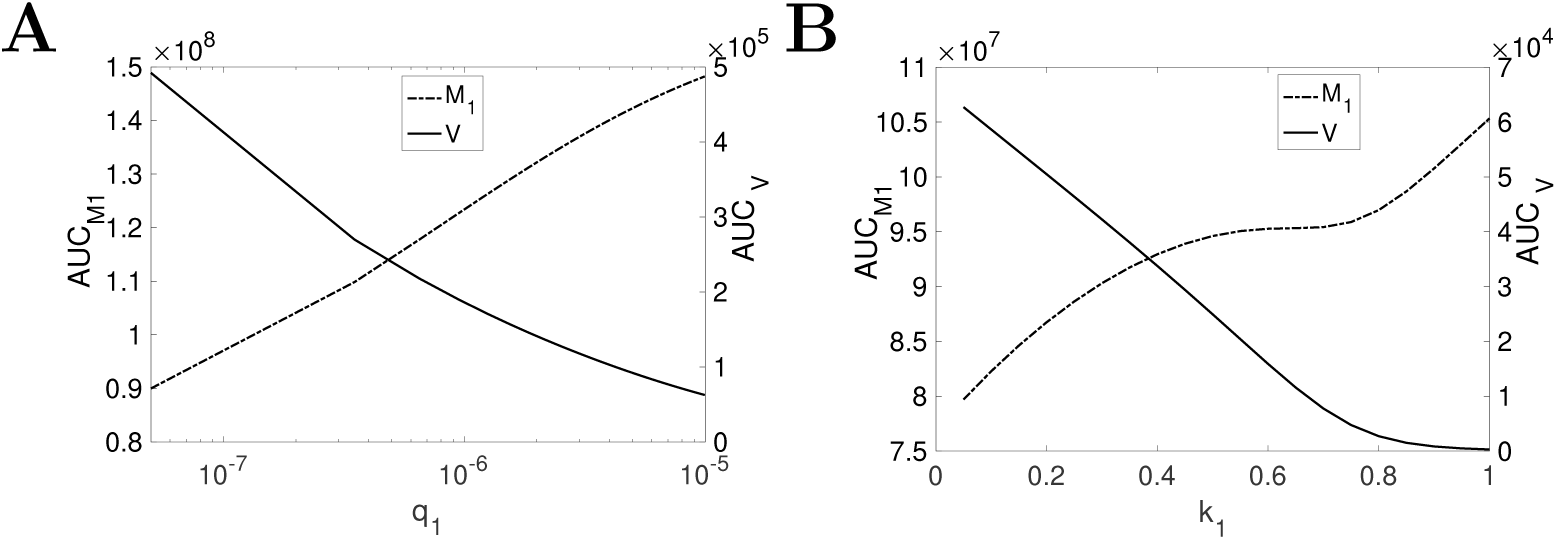
Model simulation results of AUC_*M*1_ (dashed line) and AUC_*V*_ (solid line) as functions of *q*_1_ and *k*_1_, respectively. (A) The dependence of the AUC_*M*1_ and the AUC_*V*_ on the virus-induced macrophage activation rate *q*_1_. *q*_1_ is varied from 3 *×* 10^−8^ to 1 *×* 10^−5^. (B) The change of the AUC_*M*1_ and the AUC_*V*_ to modulation of the proinflammatory macrophage activation rate *k*_1_ *∈* [0.02, 1]. The AUC_*M*1_ is indicated by dashed line, and the AUC_*V*_ is shown in solid line.

## 4. Discussion

In this work, we have studied the relationship between viral shedding and macrophage activation during influenza virus infection through numerical analysis of a mathematical model which integrates viral infection dynamics, macrophage dynamics and the essential interactions between virus and macrophages. We find based on the model that viruses with a higher ability to cause infection (e.g. a higher viral infectivity *β* or a higher production rate *p* in the model) will always lead to a higher level of viral shedding (i.e. a higher AUC_*V*_) but not necessarily a higher level of macrophage activation (i.e. a higher AUC_*M*1_). For an intermediate range of viral infectivity, the level of viral shedding and the level of macrophage activation are positively correlated (shown in Fig. 4; region II), which has been observed in avian IAV infection [32, 60, 61, 62]. But when the viral infectivity becomes sufficiently high, the level of macrophage activation declines, leading to an unexpected negative correlation with the level of viral shedding (shown in Fig. 4; region III), which is then shown to be caused by a temporary depletion of resting macrophages *M*. Our findings not only suggest that a higher viral shedding may not be accompanied by a higher macrophage response but also high-light the importance of the pool size of resting macrophages in modulating the pro-inflammatory response.

To the best of our knowledge, our model is the first work to incorporate the dynamics of heterogeneous macrophage populations into a model of influenza viral dynamics. Although no macrophage data in the literature can be used to directly test our model results, there is some indirect evidence from cytokine data to support our findings. For example, activated macrophages *M*_1_ can secrete a large amount of cytokines such as interleukin 6 (IL-6) and tumor necrosis factor (TNF) [21, 63], and evidence from experimental studies [12, 64, 65] has shown that the level of those pro-inflammatory cytokines normally rises in the early days of influenza infection and then gradually decreases afterwards—a kinetic behaviour consistent with the kinetics of *M*_1_ macrophages predicted by our model. Furthermore, alternatively activated macrophages *M*_2_ are responsible for producing anti-inflammatory cytokines, such as IL-4 and IL-10, and the time series data of IL-4 from [66] (in which mice are experimentally inoculated influenza A/PR/8/34 H1N1 virus) show qualitatively similar kinetics to that of the *M*_2_ macrophages produced by our model (i.e. an initial decrease followed by a recovery back to baseline, as shown in Fig. 3A).

Understanding the cause of symptoms due to IAV infection is an important but challenging task. Because the mechanism causing various symptoms remains unclear, we do not have the capacity to accurately model symptom dynamics. However, we can take a heuristic approach to predict the kinetics of symptoms using our model. For example, given that macrophage-mediated inflammatory response contributes to the formation of illness [24, 64, 65], then if we assume a positive correlation between symptom dynamics and the number of *M*_1_ macrophages, our model predicts a delayed presence, peak and resolution of symptoms compared to viral shedding dynamics (Fig. 3A). This prediction is qualitatively consistent with a previous finding that viral shedding preceded the occurrence of symptoms by approximately one day and finished earlier than the symptom resolution [67]. Although there are assumptions to be validated by further experiments, such as the positive correlation between the formation of symptoms and the dynamics of *M*_1_ macrophages, our model provides promising directions to probe the mechanism of symptom formation and establish the relationship between symptoms and immune response dynamics.

Further, macrophages have been shown to have pathological effects in mice infected with highly pathogenic influenza virus [68, 69]. Our model results provide new insight into the possible mechanisms for regulation of macrophage activation, suggesting that the pathological effects can be minimized by influencing the replenishment rate and reducing the available number of resting macrophages (e.g., region III Fig. 4). This may decrease macrophage accumulation and restrict the strength of inflammatory responses. For example, a study in [70] has shown that lethality of IAV infection to mice could be ameliorated when interferon-I (IFN-I) signalling is blocked. The similar knockout can be applied to macrophages and modulate the activation process of macrophages.

Our model can be extended to study other biological processes which are highly dependent upon macrophage dynamics in influenza infection. For instance, macrophages have been shown to have an important role in effective activation of adaptive immunity [12, 68], and quantifying the impact of macrophages upon adaptive immune responses in influenza infection will be a promising direction for further study. Also, biological activities of cell surface mucin (cs-mucin) glycoproteins, MUC1 particularly, have been shown to have an important role in reducing the severity of influenza infection, as reviewed in [71]. MUC1 provides two-fold protection to the host—a physical barrier to prevent virus from infecting healthy cells and more importantly a regulator of host inflammatory responses via inhibition of signalling path-ways on macrophages [72]. Our model has the capacity to model the two protective roles of MUC1. For instance, we could model the physical effect of MUC1 against influenza infection by reducing the viral infectivity *β* or model the inhibitory effect of MUC1 on inflammatory response by reducing the activation rate of *M*_1_ macrophages. With the data available from [72], we may be able to quantitatively study the effect of MUC1 on reduction of infection severity and inflammatory responses in influenza infection. Another potential application of our model is to predict the effect of novel antiviral treatments, such as Pam2Cys, a novel immunomodulator shown to be able to enhance protection against influenza in mice by stimulating innate immunity and recruiting macrophages to the site of infection [73, 74]. These applications are beyond the scope of this paper and are left for future work.

## Author contributions

**Ke Li:** Conceptualization, Methodology, Software, Formal analysis, Writing-Original Draft. **James M. McCaw** Methodology, Formal analysis, Writing-Review and Editing, Supervision. **Pengxing Cao:** Methodology, Formal analysis, Writing-Review and Editing, Supervision

### Acknowledgements

Ke Li is supported by a Melbourne Research Scholarship. This work was supported by an Australian Research Council (ARC) Discovery Project (DP170103076) and a National Health and Medical Research Council (NHMRC) funded Centre for Research Excellence in Infectious Diseases Modelling to Inform Public Health Policy (1078068).

