## Supplementary Materials 1 for "Modelling within-host macrophage dynamics in influenza virus infection"

**Ke Li, James M. McCaw, Pengxing Cao**

**The parameters values in Table 1 in the main text, otherwise specified, are used for following simulation.**

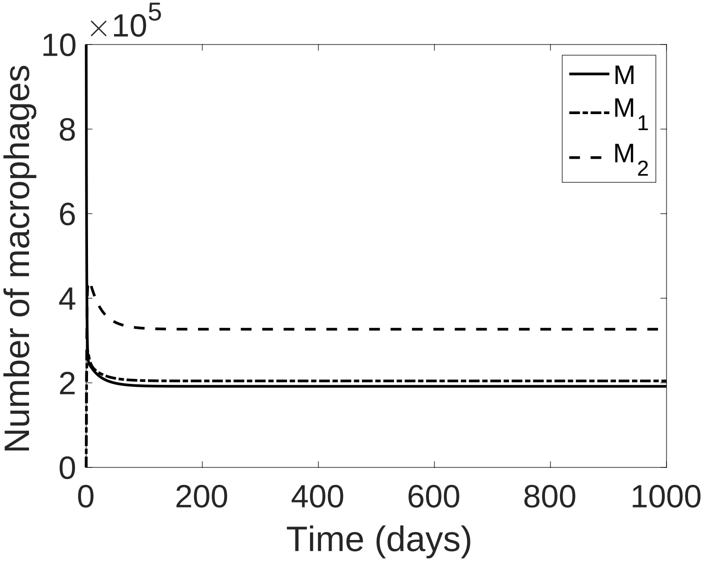


Figure 1: Model simulation results of Eq. 1–Eq. 3 in the main text, which give macrophage dynamics in the absence of viral infection. The solid line denotes resting inactivated macrophages $M$; the dot-dashed line denotes classically activated macrophages $M_1$, and the dot line denotes alternatively activated macrophages $M_2$.

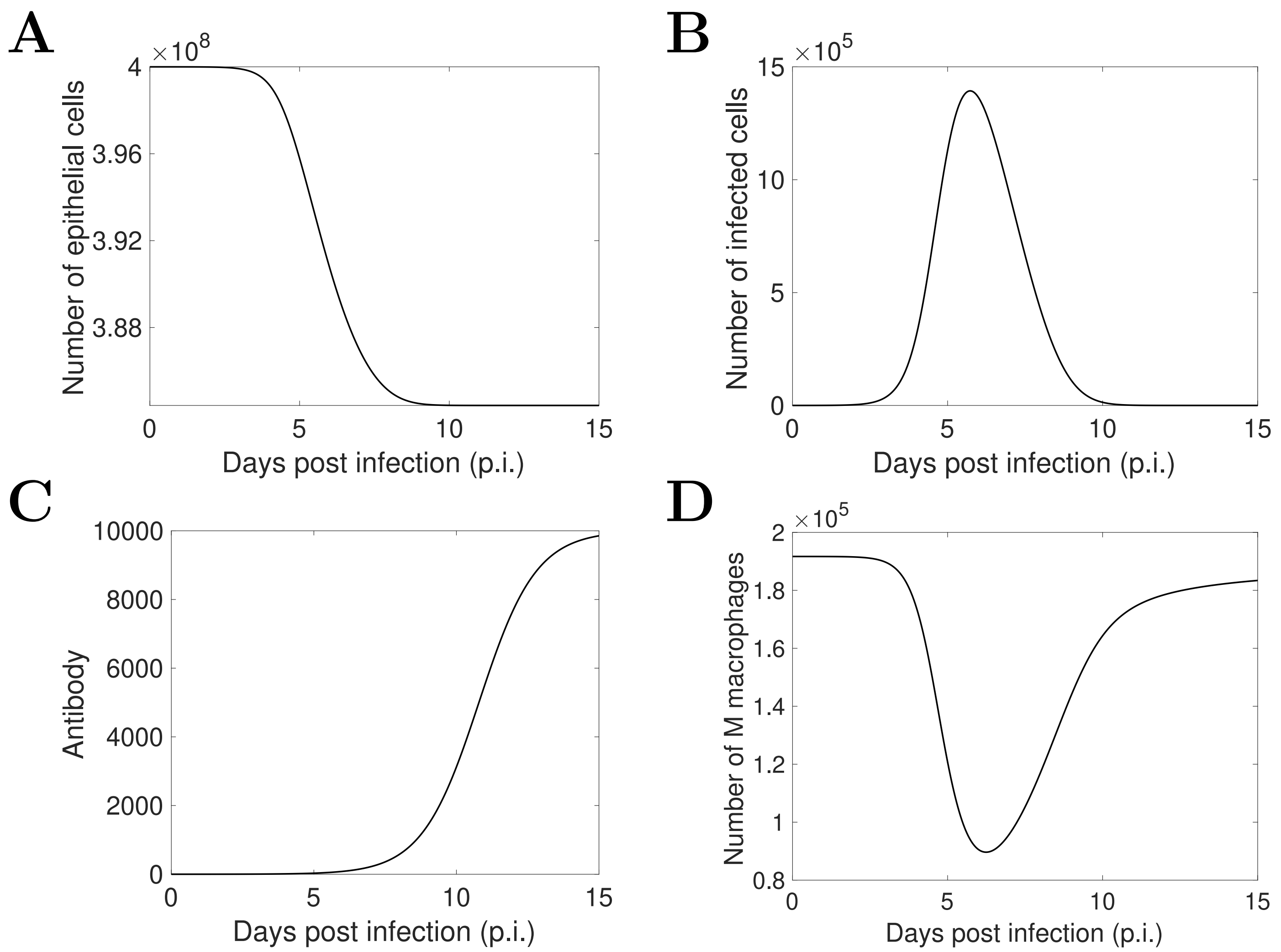


Figure 2: Model simulation results (A–D) of Eq. 7 (epithelial cells), Eq. 8 (infected cells), Eq. 10 (antibodies) and Eq. 4 (resting macrophages $M$).

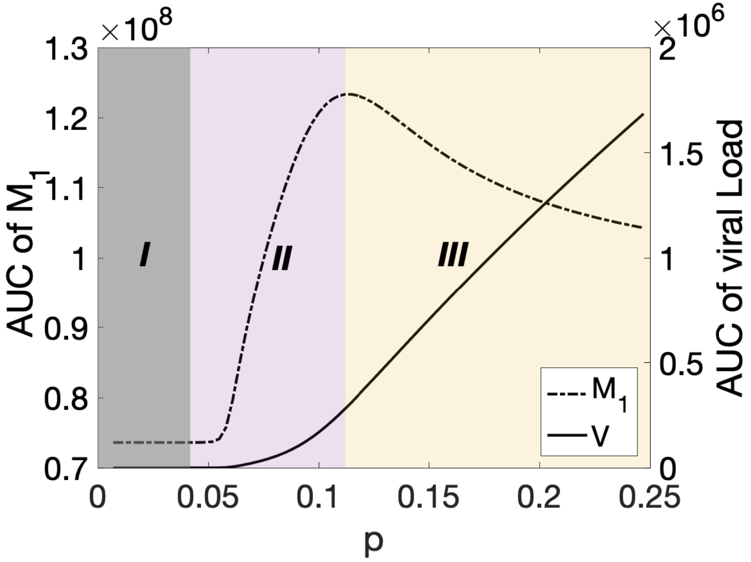


Figure 3: Model simulation results of the change of the $\mathrm{AUC}_{M1}$ (dashed line) and the $\mathrm{AUC}_V$ (solid line) to modulation of viral replication rate $p \in [7.1 \times 10^{-3}, 2.5 \times 10^{-1}]$. See the main text for detailed method of separation of regions I, II and III.

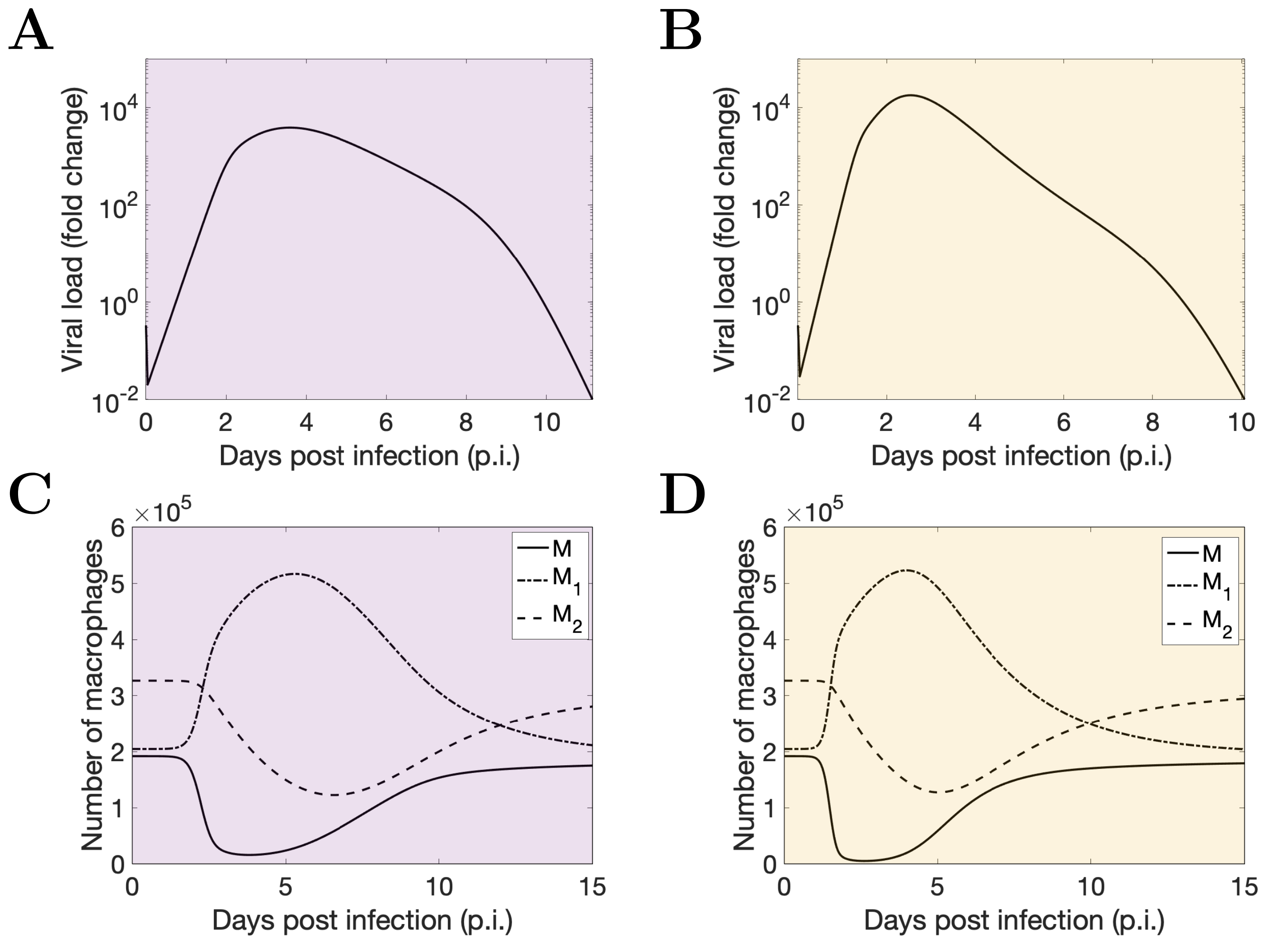


Figure 4: **Dependence of model behaviours on different viral replication rate.** (A) and (B) are dynamics of viral shedding in regions II ($p = 10 \times 10^{-2}$) and III ($p = 2 \times 10^{-1}$), respectively. (D) and (E) show macrophage dynamics in regions II and III (the same $p$ accordingly).

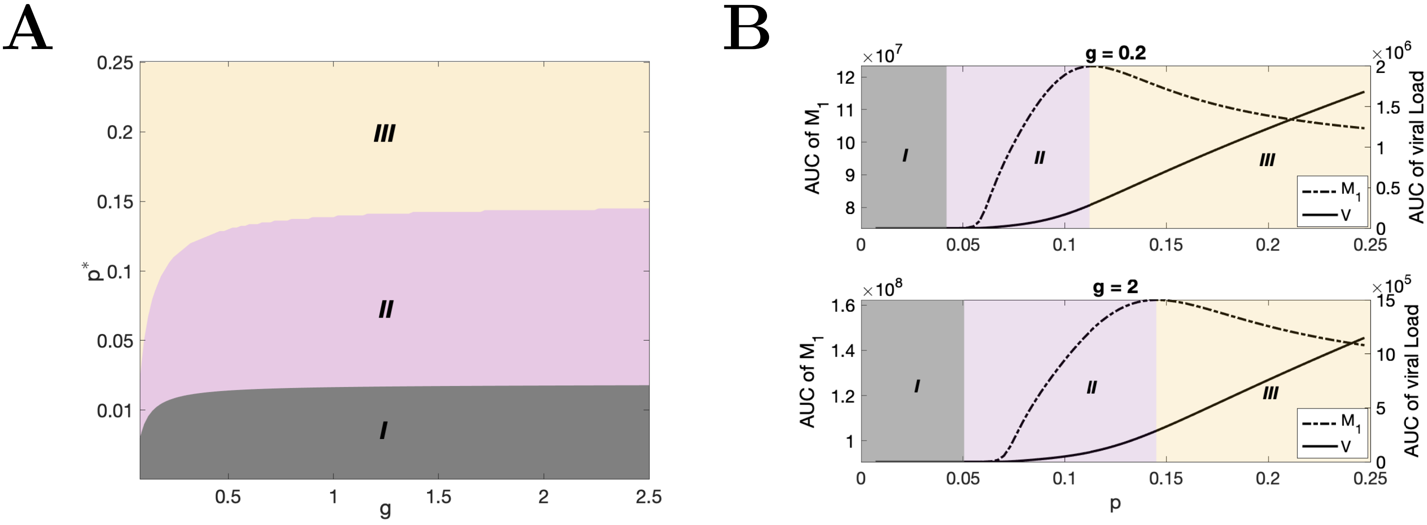


Figure 5: **Model simulation results of the $\text{AUC}_{M1}$ and $\text{AUC}_V$ as functions of of $M$ regrowth rate $g$.** (A) shows how the critical $p$ that divides regions I, II and III changes as $g$ increases. (B) The change of the $\text{AUC}_{M1}$ and the $\text{AUC}_V$ to modulation of the viral replication rate $p \in [7.1 \times 10^{-3}, 2.5 \times 10^{-1}]$ with $g = 0.2$ and $g = 2$, respectively.

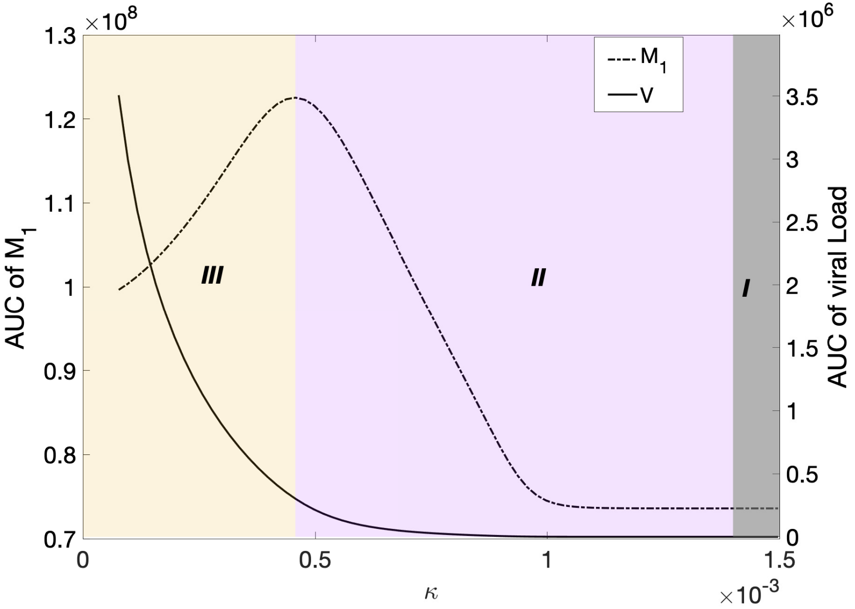


Figure 6: Model simulation results of the change of the $\mathrm{AUC}_{M1}$ (dashed line) and the $\mathrm{AUC}_V$ (solid line) to modulation of the rate of $M_1$ macrophages engulf virus $\kappa \in [7.7 \times 10^{-5}, 1.5 \times 10^{-3}]$. See the main text for detailed method of separation of regions I, II and III.

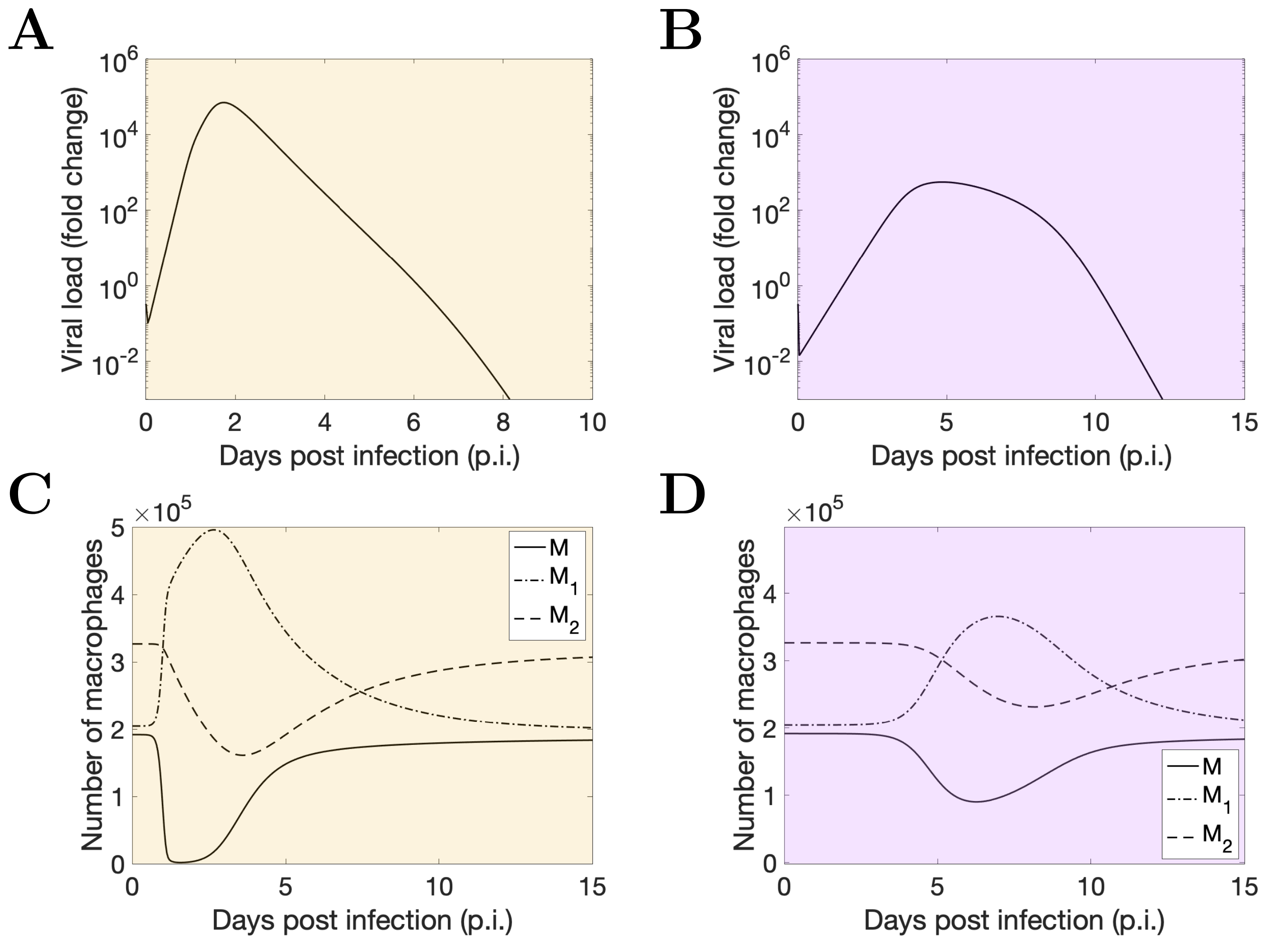


Figure 7: **Dependence of model behaviours on the rate of $M_1$ macrophages engulf virus.** (A) and (B) are dynamics of viral shedding in regions III ($\kappa = 3 \times 10^{-4}$) and II ($\kappa = 4.6 \times 10^{-4}$), respectively. (C) and (D) show macrophage dynamics in regions III and II (the same $\kappa$ accordingly).

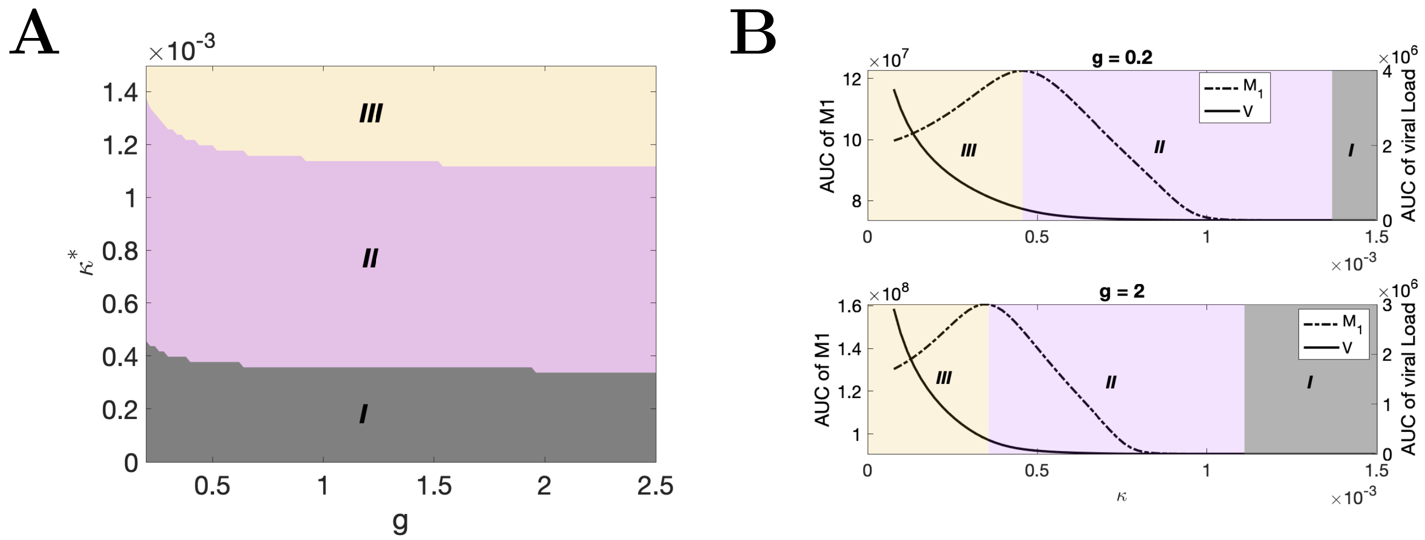


Figure 8: **Model simulation results of the $\text{AUC}_{M1}$ (dashed line) and $\text{AUC}_V$ (solid line) as functions of $M$ regrowth rate $g$.** (A) shows how the critical $\kappa$ that divides regions I, II and III changes as $g$ increases. (B) The change of the $\text{AUC}_{M1}$ and the $\text{AUC}_V$ to modulation of the engulfment rate of virus by $M_1$, $\kappa \in [7.7 \times 10^{-5}, 1.5 \times 10^{-3}]$ with $g = 0.2$ and $g = 2$, respectively.
