## Supplementary Materials 2 for "Modelling within-host macrophage dynamics in influenza virus infection"

**Ke Li, James M. McCaw, Pengxing Cao**

The mathematical calculation below shows that the equilibrium of macrophages (Fig. 6A in the main text) will gradually approach to the maximal values when the regrowth rate of macrophages $M$, $g$, goes sufficiently large.

Given Eq. 4 and $dM/dt = 0$ at equilibrium at any given time, we have
$M_0 - (1 + \alpha_1(t) + \alpha_2(t))M(t) = \Gamma(t)/g$,

where

$$\alpha_1(t) = \frac{k_1(M_2(t)) + q_1 I(t) + q_2 V(t)}{k_{-1} + \delta},$$
$$\alpha_2(t) = \frac{k_2(M_1(t))}{k_{-2} + \delta},$$
$$\Gamma(t) = (k_1(M_2(t)) + k_2(M_1(t)) + q_1 I(t) + q_2 V(t) - k_{-1} - k_{-2})\, M_0,$$

and

$$k_1(M_2(t)) = k_1 / \left(1 + s_1 \frac{M_2(t)}{M_0}\right),$$
$$k_2(M_1(t)) = k_2 \left(1 + s_2 \frac{M_1(t)}{M_0}\right).$$

Then, the maximal number of $M(t)$ at any unit time is

$$M^*(t) = \frac{M_0}{1 + \alpha_1(t) + \alpha_2(t)}, \quad \text{as} \quad g \to +\infty$$
